# Discovery of a two protease DNA damage checkpoint recovery mechanism

**DOI:** 10.1101/303297

**Authors:** Peter E. Burby, Zackary W. Simmons, Jeremy W. Schroedert, Lyle A. Simmons

## Abstract

The DNA damage response is a signaling pathway found throughout biology. In many bacteria the DNA damage checkpoint is enforced by inducing expression of a small, membrane bound inhibitor that delays cell division providing time to repair damaged chromosomes. How cells sense successful DNA repair and promote checkpoint recovery is unknown. By using a high-throughput, forward genetic screen, we identified two unrelated proteases, YlbL and CtpA, that promote DNA damage checkpoint recovery in *Bacillus subtilis*. Deletion of both proteases leads to accumulation of the checkpoint protein YneA. DNA damage sensitivity and increased cell elongation in protease mutants depends on *yneA*. Further, expression of YneA in protease mutants was sufficient to inhibit cell proliferation. Finally, we show that one of the two proteases, CtpA, directly cleaves YneA *in vitro*. With these results, we report the mechanism for DNA damage checkpoint recovery in bacteria that use membrane bound cell division inhibitors.

## Introduction

The <u>D</u>NA <u>d</u>amage <u>r</u>esponse (DDR, SOS response in bacteria) is an important pathway for maintaining genome integrity in all domains of life. Misregulation of the DDR in humans can result in various disease conditions (1, 2), and in bacteria the SOS response has been found to be important for survival under many stressors (3–5). The DNA damage response in all organisms results in three principle outcomes: a transcriptional response, DNA repair, and activation of a DNA damage checkpoint (6–8). In eukaryotes, the G1/S and G2/M checkpoints are established by checkpoint kinases, which transduce the signal of DNA damage through inactivation of the phosphatase Cdc25 (7). Checkpoint kinase dependent inhibition of Cdc25 leads to accumulation of phosphorylated cyclin dependent kinases, which prevents cell cycle progression (7). In bacteria, the SOS-dependent DNA damage checkpoint relies on expression of a cell division inhibitor, though the type of inhibitor varies between bacterial species.

In *Escherichia coli*, the SOS-dependent DNA damage checkpoint is the best understood bacterial checkpoint (6). Upon activation of the SOS response, the cytoplasmic cell division inhibitor SulA is expressed (9). SulA accumulation leads to a block in septation and assembly of the cytokinetic ring by FtsZ, a homolog of eukaryotic tubulin (10, 11). SulA binds directly to FtsZ (12) and inhibits FtsZ polymerization (13, 14). Recovery from the SulA-induced checkpoint occurs through proteolysis of SulA. Lon is the primary protease responsible for clearing SulA (15–17), although ClpYQ (HslUV) were found to contribute to SulA degradation in the absence of Lon (18–20). Thus, the mechanisms of DNA damage checkpoint activation by the cytoplasmic protein SulA and subsequent recovery are well understood in *E. coli*. The SulA-dependent checkpoint, however, is restricted to *E. coli* and a subset of closely related bacteria. It is becoming increasingly clear that most other bacteria use a DNA damage checkpoint with an entirely different mechanism of enforcement and recovery.

An evolutionarily broad group of bacterial organisms have been shown to use a notably different mechanism as a DNA damage checkpoint (21–24). In these Gram-positive and Gram-negative organisms, a small protein with a transmembrane domain is expressed that inhibits cell division without targeting FtsZ. One example is in the Gram-negative bacterium *Caulobacter crescentus*, where the SidA and DidA proteins bind to the essential membrane bound divisome components, FtsW/N that contribute to peptidoglycan remodeling (21, 25). Another example is the Gram-positive bacterium *Bacillus subtilis* in which the SOS-dependent cell division inhibitor is YneA (22). YneA contains an N-terminal transmembrane domain with the majority of the protein found in the extracellular space (26). Upon SOS activation, LexA-dependent repression of *yneA* is relieved and *yneA* is expressed (22). Increased expression of *yneA* results in cell elongation, though FtsZ ring formation still occurs (26), suggesting YneA inhibits cell division through a mechanism distinct from that of SulA. Further investigation found that overexpressed YneA is released into the medium, and that full length YneA is likely the active form of the protein (26). The mechanism(s) responsible for YneA inactivation is unknown. Therefore, although the use of a small, membrane bound cell division inhibitor is wide-spread among bacteria, in all cases studied the mechanism of checkpoint recovery remains unknown(21–25).

We report a set of forward genetic screens to three different classes of DNA damaging agents using transposon mutagenesis followed by deep sequencing (Tn-seq). Our screen identified two proteases, YlbL and CtpA, that are important for growth in the presence of DNA damage. Mechanistic investigation demonstrates that YlbL and CtpA have overlapping functions, and in the absence of these two proteases, DNA damage-dependent cell elongation is increased and checkpoint recovery is slowed. A proteomic analysis identified accumulation of YneA in the double protease mutant. We also found that DNA damage sensitivity of protease mutants depends solely on *yneA*. Further, we show that CtpA is able to digest YneA in a purified system. With these results, we present a model of DNA damage checkpoint recovery for bacteria that use the more wide-spread mechanism employing a small, membrane bound cell division inhibitor.

## Results

### Forward genetic screen rationale and analysis

In order to better understand the DNA damage response in bacteria, we performed three forward genetic screens using *B. subtilis*. We generated a transposon insertion library consisting of more than 120,000 distinct insertions (Table S1). The coverage of each transposon mutant in the library was plotted against the genome coordinates, which showed that the distribution of insertions was approximately uniform across the chromosome in the population of mutants (Fig 1A). Two small exceptions were detected where coverage decreased. Decreased coverage corresponds to regions where many essential genes are clustered (Fig 1A, arrow heads). To identify mutants important for the DNA damage response, we grew parallel cultures of either control or DNA damage treatment over three growth periods, modelling our experimental design after a previous report (27; Fig 1B). Mitomycin C (MMC), methyl methane sulfonate (MMS), and phleomycin (Phleo) were chosen for screening because these agents represent three different classes of antibiotics that damage DNA directly. MMC causes inter- and intra-strand crosslinks and larger adducts (28, 29), MMS causes smaller adducts consisting of DNA methylation (30), and Phleo results in single and double stranded breaks (31, 32). As a result, we reasoned that the combined data would provide a collection of genes that are generally important for the DNA damage response.

**Figure 1.**
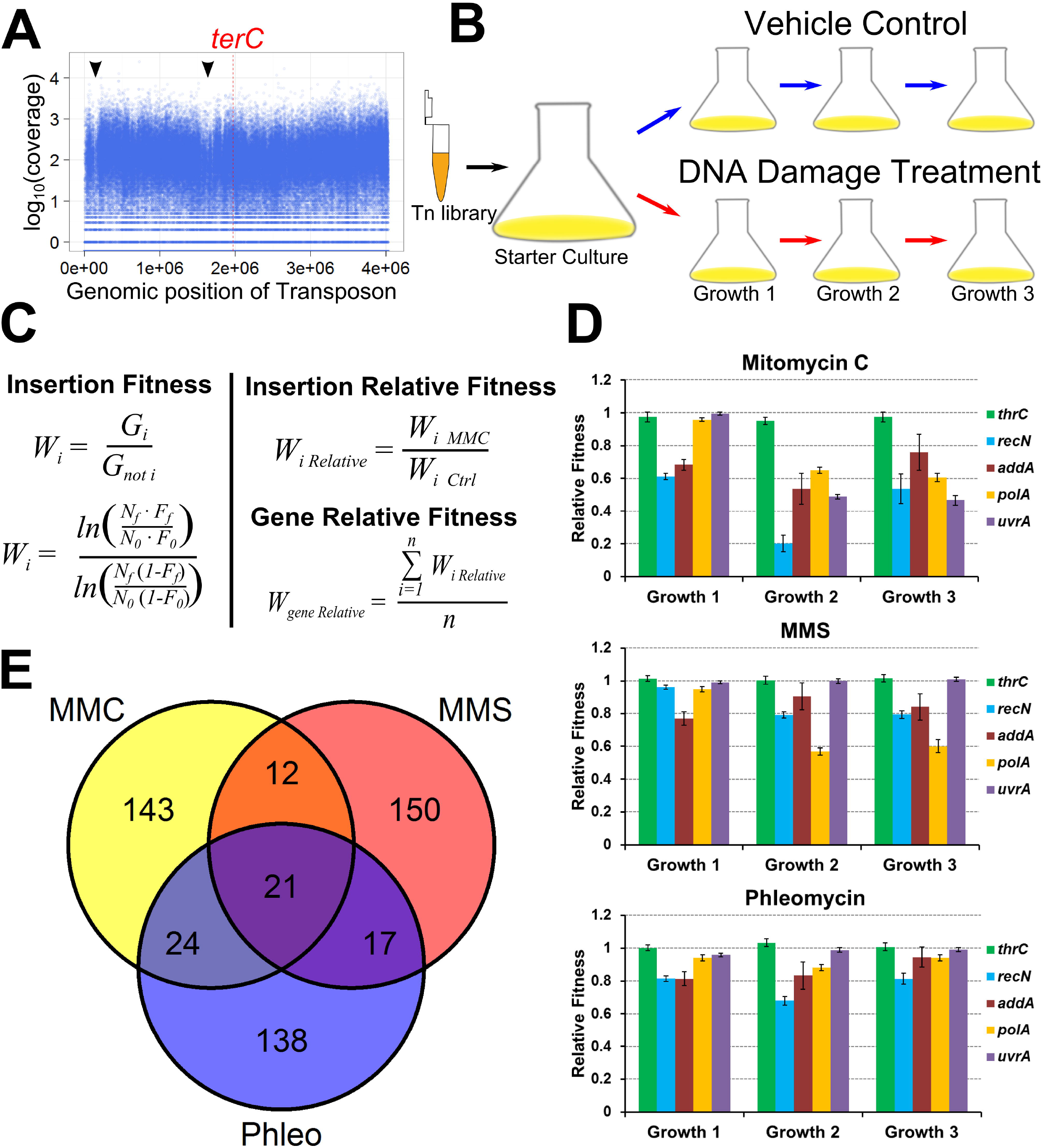
Forward genetic screen experimental design and data analysis. **(A)** A plot of the log10 of insertion coverage on the y-axis and genomic position in nucleotides on the x-axis. **(B)** Experimental design for Tn-seq experiments. The transposon library was used to inoculate starter cultures to allow cultures to reach exponential phase. Cultures were split into control and treatment and grown for three growth periods. **(C)** Equations used to calculate relative fitness (W, fitness; G, generations, N, number of cells at the start (N_0_) or end (N_f_) of growth period; F, insertion frequency at the start (F_0_) or end (F_f_) of growth period; n, number of insertions used to calculate average). **(D)** The mean gene relative fitness is plotted as a bar graph for the genes indicated for all three Tn-seq experiments, error bars represent the 95% confidence interval. **(E)** A Venn diagram depicting overlaps of the 200 genes with the lowest fitness and an adjusted p-value less than 0.01 for all three Tn-seq experiments in growth period two.

After sequencing, we performed quality control analysis. First, given that sequencing data are count data, the distribution of the coverage should be log-normal (33). Indeed, the distribution of each replicate for the initial library and starter culture samples is approximately log-normal (Fig S1A). We also found that the distributions for the remaining time points of the pooled replicates followed an approximate log-normal distribution (Fig S1B). The sequencing data and viable cell count data (Table S1) were used to calculate the fitness of each insertion mutant in each condition (Fig 1C; 34, 35). The relative fitness of each insertion was calculated by taking the ratio of treatment to control (Fig 1C), thereby isolating fitness effects of the treatments. The relative fitness of each gene was determined by averaging the relative fitness calculated for each insertion within a gene (Fig 1C). To verify that a t-test would be appropriate for determining relative fitness deviating significantly from one, we plotted the distribution of insertion relative fitness. All the distributions were normal with a mean close to one (Fig S1C). We determined the relative fitness for every gene with sufficient data (see STAR methods), and report the relative fitness values and the adjusted p-values (36) in Table S2.

### Tn-seq identified genes involved in DNA repair and genes of unknown function

Initial inspection revealed that several genes known to be involved in DNA repair (*recA, ruvAB, recN*, and *recOR* (37)) had decreased relative fitness in growth period one of all experiments (Table S2). A closer analysis of *recN, addA*, and *polA*, three genes that are found toward the top of the lists in all treatments, showed that relative fitness is less than one in most cases, though in the Phleo experiment, it appears that the cultures were adapting to the treatment by growth period three (Fig 1D). For comparison, we also plotted the relative fitness of *thrC*, a gene involved in threonine biosynthesis, and found the relative fitness to be about one in all conditions examined (Fig 1D). Importantly, insertion in *uvrA*, a component of the nucleotide excision repair machinery (37, 38), which helps repair MMC adducts but not MMS or Phleo related damage (39, 40), decreased relative fitness in growth periods 2 and 3 with MMC, but did not significantly decrease relative fitness in MMS or Phleo (Fig 1D and Table S2). Taken together, these results validate the approach by demonstrating that we were able to identify genes known to be involved in DNA repair. To identify genes required generally as part of the DNA damage response, we examined the 200 genes from growth period two with the lowest relative fitness and an adjusted p-value less than 0.01 from all three experiments (Table S3). We found that 21 genes overlapped for all three experiments (Fig 1E), some of which are known to be involved in DNA repair (*recN, addB, polA, radA*), while several genes have no known function (e.g., *ylbL* and *ctpA*) (Table S3).

### YlbL and CtpA require putative catalytic residues for function

Among the genes important for growth in the presence of DNA damage, we focused on two putative proteases YlbL and CtpA. YlbL is predicted to have three domains: a transmembrane domain, a Lon protease domain, and a PDZ domain (Fig 2A). CtpA is predicted to have four domains: a transmembrane domain, a S41 peptidase domain, a PDZ domain, and a C-terminal peptidoglycan (PG) binding domain (Fig 2A). In all three Tn-seq experiments, the relative fitness of insertions in either *ylbL* or *ctpA* was significantly less than one in the second and third growth periods (Fig 2B), suggesting that absence of either protease results in sensitivity to DNA damage. In contrast, a control gene *amyE*, which is involved in starch utilization, had a relative fitness of approximately one in all conditions examined (Fig 2B). To verify the Tn-seq results, we constructed clean deletions of *ylbL* and *ctpA* and found both mutants to be sensitive to DNA damage in a spot-titer assay (Fig 2C). Each phenotype was also complemented by ectopic expression of each protease in its respective mutant background (Fig 2C). To identify putative catalytic residues, we aligned the protease domain of YlbL to LonA and LonB from *B. subtilis* and Lon from *E. coli*. The sequence alignment revealed that YlbL contains a putative catalytic dyad consisting of a serine (S234) and a lysine (K279) (Fig S2A). Similarly, we aligned CtpA to its homologs CtpB from *B. subtilis* and Prc from *E. coli*, which identified a putative catalytic triad consisting of a serine (S297), a lysine (K322), and a glutamine (Q326) (Fig S2B). To test whether these putative catalytic residues were required for function, we attempted to complement the DNA damage sensitivity phenotype via ectopic expression of serine and lysine mutants. Both the serine and lysine mutants of YlbL and CtpA failed to complement the deletion phenotypes (Fig 2C). The variants and the wild-type proteases were ectopically expressed to the same level *in vivo* (Fig 2D), suggesting that the lack of complementation is not due to instability caused by the amino acid changes. With these results, we conclude that protease activity is required for YlbL and CtpA to function in response to DNA damage.

**Figure 2.**
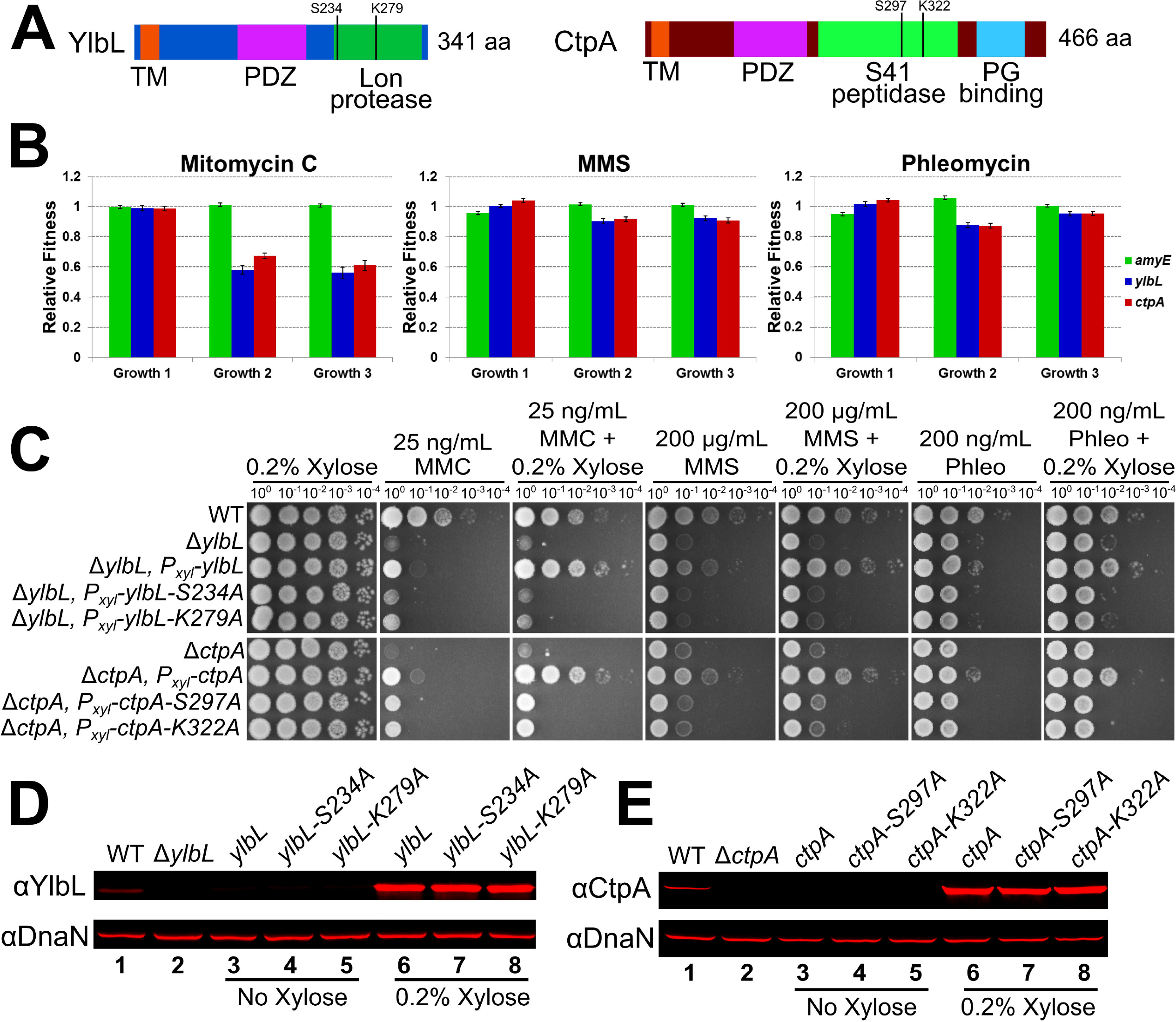
YlbL and CtpA require putative catalytic residues for function. **(A)** Schematics of YlbL and CtpA proteins depicting the domain organization. **(B)** The mean gene relative fitness is plotted as a bar graph for *amyE, ylbL*, and *ctpA* for all three Tn-seq experiments, error bars represent the 95% confidence interval. **(C)** Spot titer assays using the genotypes indicated and plated on LB agar media containing the indicated drugs. **(D)** Western blot analysis of cell lysates from the indicated genotypes grown with or without xylose using antiserum against YlbL (upper panel) and DnaN (lower panel). **(E)** Western blot analysis of cell lysates from the indicated genotypes grown with or without xylose using antiserum against CtpA (upper panel) and DnaN (lower panel).

### *ylbK* disruption results in a polar effect on *ylbL*

We noticed that *ylbK*, the gene upstream of *ylbL*, had a phenotype similar to *ylbL* in the Tn-seq experiments (Table S2 & S3). We tested whether *ylbKL* functioned together in the DNA damage response. Deletion of *ylbK* resulted in sensitivity to MMC (Fig S3A). Ectopic expression of *ylbK* failed to complement the Δ*ylbK* phenotype (Fig S3A). Given that *ylbK* is upstream of *ylbL* we attempted to complement the Δ*ylbK* phenotype using *ylbL* and found that sensitivity to MMC was rescued (Fig S3A). Closer examination of the *ylbKL* locus revealed that a putative ribosome binding site (RBS) for *ylbL* translation was present within the 3’ end of *ylbK* (Fig S3B). Thus, a second deletion of *ylbK* was made (Δ*ylbK-2*) which included deletion of the codons for all but the first 3 and the last 14 amino acids, leaving the RBS for *ylbL* intact (Fig S3B). This deletion was not sensitive to MMC (Fig S3A). Western blotting revealed that the initial Δ*ylbK* construct did not express YlbL, whereas Δ*ylbK-2* did (Fig S3C). As a result, we conclude that disruption of *ylbK* results in a polar effect on *ylbL*, indicating that YlbL functions independently of YlbK.

In order to better understand the prevalence of false positives in Tn-seq experiments we attempted to validate the MMC phenotypes of the forty genes with the lowest relative fitness values in the second growth period of the MMC experiment. Intriguingly, we found that seven additional genes, *queA, ylmG, lgt, ylmE, sdaAB, cymR*, and *ywrC*, resulted in no sensitivity to MMC when deleted (Table 1). We also found that the genomic loci of *queA, ylmG, ylmE*, and *sdaAB*, were proximal genes with validated phenotypes (Table 1). Taken together, our results underscore the importance of validating results from forward genetic screens.

**Table 1.**
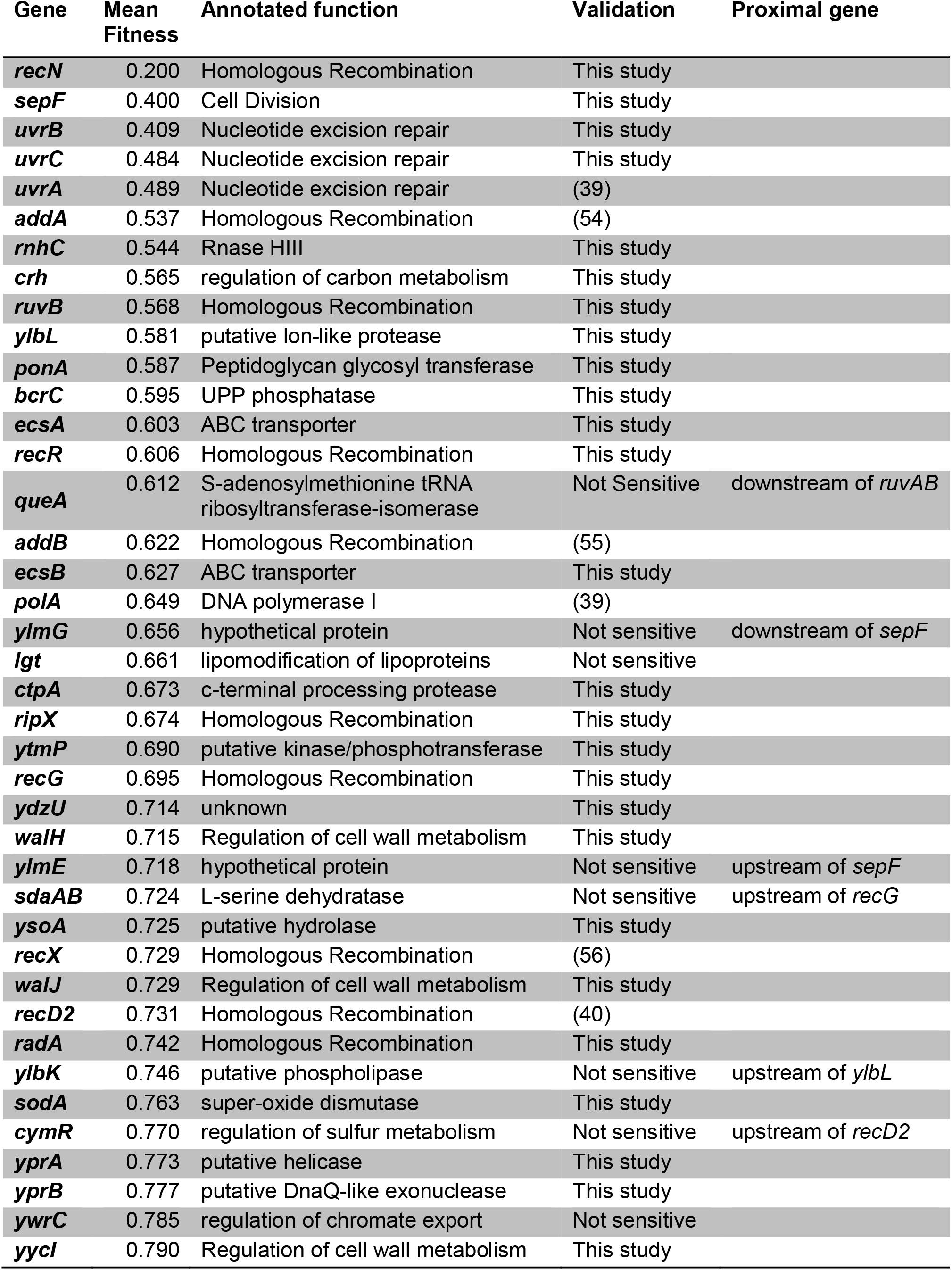
Tn-seq yields many false positive results. The forty genes with the lowest relative fitness in the second growth period of MMC Tn-seq experiment are listed. Each gene was deleted and the deletion mutants were tested for sensitivity to MMC using a spot titer assay and a range of MMC concentrations. Genes labeled as not sensitive had no difference in growth relative to the WT strain on MMC containing media, with the exception of *ylbK*, which resulted in a polar effect on *ylbL* (see figure S3).

### YlbL and CtpA have overlapping functions

The similarity in phenotypes led us to hypothesize that YlbL and CtpA have overlapping functions. To test this we performed a cross-complementation experiment using spot-titer assays for MMC sensitivity. Over-expression of YlbL, but not YlbL-S234A, complemented a *ctpA* deletion (Fig 3A & B). Similarly, over-expression of CtpA, but not CtpA-S297A, complemented a *ylbL* deletion (Fig 3A & B). In addition, deletion of both proteases rendered *B. subtilis* hypersensitive to MMC, even more so than loss of *uvrA*, which codes for the protein responsible for recognizing MMC adducts as part of nucleotide excision repair (Fig 3C; 37, 38). To further test the hypothesis that YlbL and CtpA have overlapping functions, we over-expressed each of the proteases separately in the double protease mutant background and observed a complete rescue of MMC sensitivity upon expression of the wild type (WT), but not the serine variants (Fig 3D & E).

**Figure 3.**
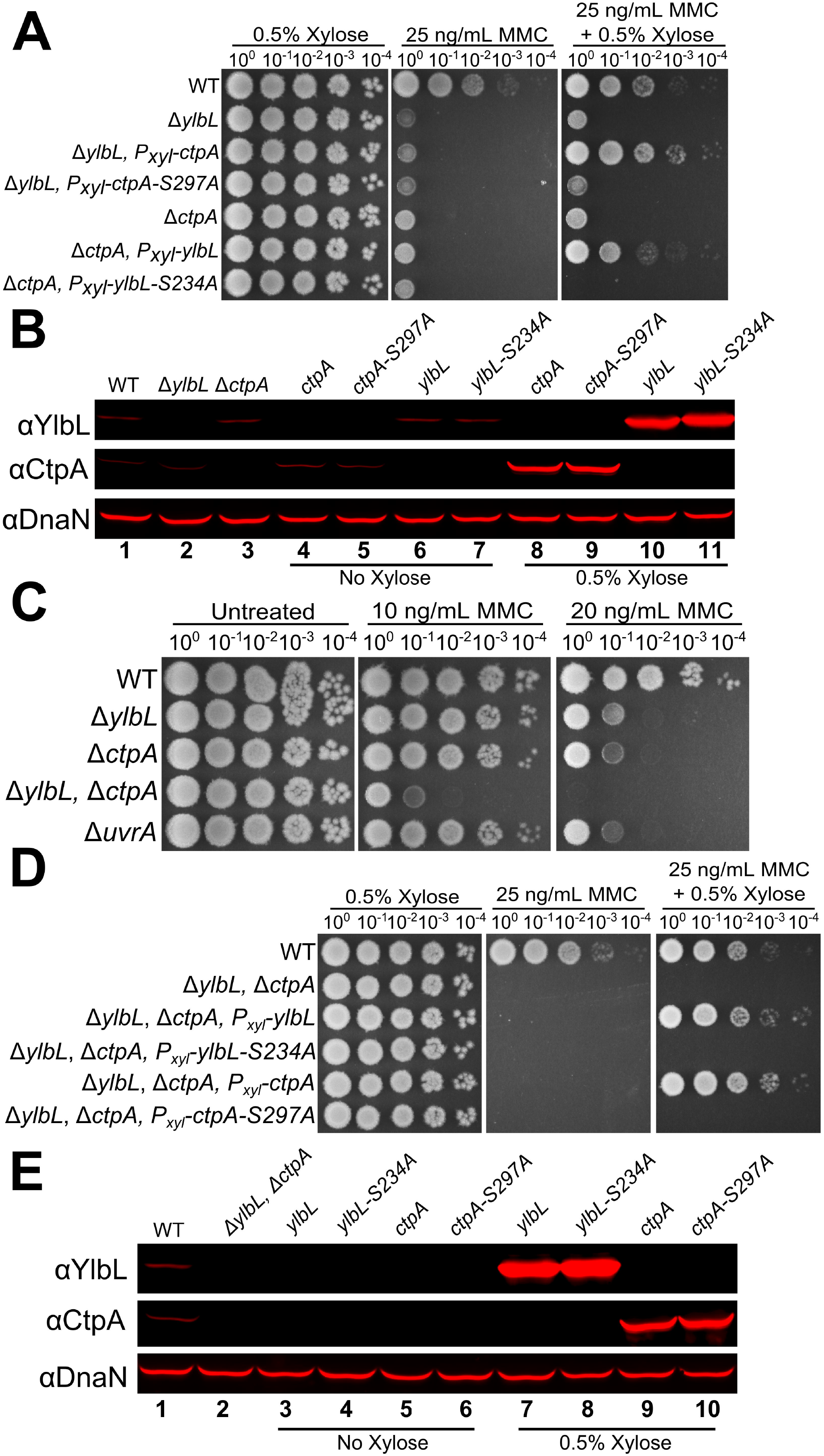
YlbL and CtpA have overlapping functions. **(A)** Spot titer assay using the indicated genotypes and media. **(B)** Western blot analysis of cell lysates from the genotypes in panel A, using the indicated antiserum. (**C & D**) Spot titer assay using the indicated genotypes and media. **(E)** Western blot analysis of cell lysates from the genotypes indicated in panel D, using the indicated antiserum.

### DNA damage-dependent cell division delay is increased in protease deletions

The experiments performed thus far cannot distinguish between sensitivity to MMC resulting from cell death, growth inhibition or both. To determine whether sensitivity arises from cell death, we performed a survival assay using an acute treatment of MMC. We detected a slight decrease in percent survival as MMC concentration increased in the Δ*ylbL* and the double mutant strain (Fig S4A). We compared the decrease in percent survival in single and double protease mutants to a Δ*uvrA* strain, which has been shown previously to be acutely sensitive to MMC (39). The strain lacking *uvrA* was very sensitive to an acute treatment of MMC (Fig S4A), whereas, the double protease deletion strain was significantly less sensitive to acute exposure compared with Δ*uvrA* (compare Fig 3C & S4A). Taken together, we conclude that MMC sensitivity of the protease mutants observed in spot-titer assays is primarily caused by growth inhibition.

We hypothesized that sensitivity to DNA damage resulting from growth inhibition could also be explained by inhibiting cell proliferation, or inhibiting cell division rather than cell growth. To distinguish between these two possibilities, we measured cell length, because inhibition of proliferation should be observed as an increase in cell length, consistent with a failure in checkpoint recovery. Thus, we designed a MMC recovery assay, reasoning that following treatment with MMC, cells lacking YlbL, CtpA, or both, would remain elongated showing slower checkpoint recovery relative to the WT strain. We grew cultures either in a vehicle control or in the presence of MMC. After a two hour treatment, the MMC containing media was removed and cells were washed. Cells were then transferred to fresh media without MMC and allowed to continue growing to assay for checkpoint recovery. Although cells appeared to be elongated in the Δ*ylbL* and double mutant strains, there was heterogeneity in the population (Fig 4A). As a result, we measured the cell length of at least 900 cells for each genotype and each condition and plotted the cell length distributions as histograms (Fig 4B). There was no difference in the vehicle control cell length distributions (Fig 4B). The MMC treatment of all strains resulted in a rightward shift in the distribution for all strains (Fig 4B, compare upper panels). When comparing the protease deletions to the WT, the difference in distribution could be visualized by considering the percentage of cells greater than 6.75 μm in length, which is about three cell lengths of 2.25 μm each. We found that deletion of *ylbL* resulted in an increase in the percentage of cells longer than 6.75 μm in MMC treated cultures and after both 2 hours and 4 hours of recovery (Fig 4C). Deletion of *ctpA*, however, resulted in a very slight, though significant (p-value = 0.0142 for one-tailed Z-test), increased percentage of cells longer than 6.75 μm after 4 hours of recovery (Fig 4C). The double mutant resulted in a percentage of cells slightly greater than Δ*ylbL* alone after both 2 hours (p-value = 0.0001 for one-tailed Z-test) and 4 hours (p-value = 0.0088 for one-tailed Z-test) of recovery (Fig 4C).

**Figure 4.**
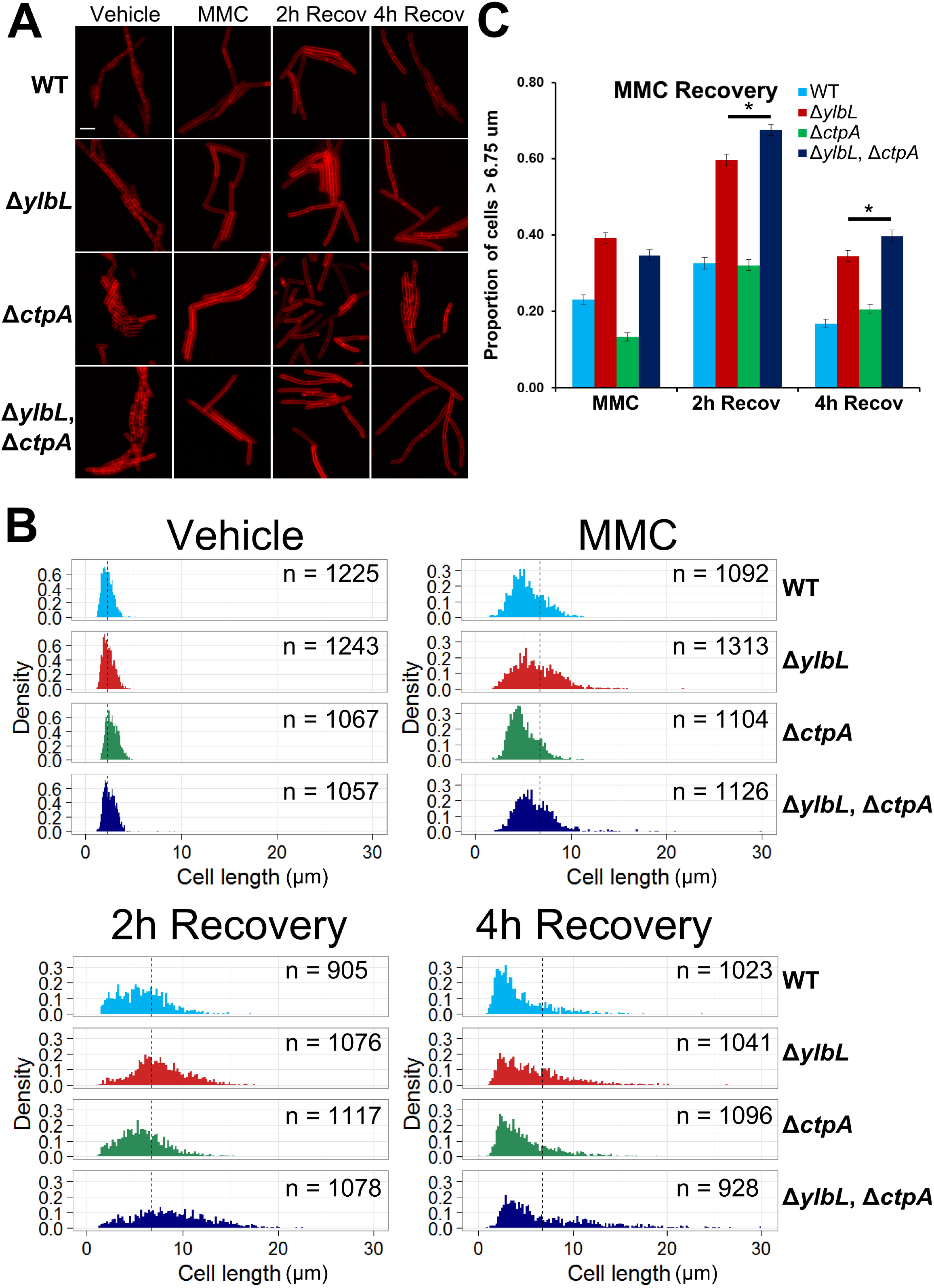
DNA damage delays cytokinesis in cells with protease deletions. **(A)** Representative micrographs of cells with the indicated genotypes at the indicated time points. Membranes were stained with FM4-64. Scale bar is 5 μm. **(B)** Cell length distributions plotted as histograms. The y-axis in all graphs is normalized by the sample size yielding the density, and the x-axis is the cell length in μm. The number of cells scored in each distribution is indicated as “n=” and the genotype of each strain is indicated. The dotted vertical line in the “Vehicle” distributions is plotted at 2.25 μm, the approximate mean for all strains. The dotted vertical line in the remaining distributions is at 6.75 μm, which is three times the average length of untreated cells. **(C)** The proportion of cells represented by the histograms in panel B with length greater than 6.75 μm is plotted as a bar graph. The error bars represent the standard deviation, 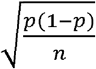, where p represents the proportion and n is the sample size). The asterisk indicates a p-value less than 0.05.

Taken together, we conclude that YlbL is the primary protease under these conditions, with CtpA also contributing. We also conclude that cells lacking YlbL or both YlbL and CtpA take longer to divide following exposure to MMC, which is consistent with DNA damage sensitivity resulting from inhibition of cell proliferation. Further, the observation of inhibition of cell proliferation suggests that YlbL and CtpA proteases could be important for DNA damage checkpoint recovery (see below).

### YlbL and CtpA levels are not regulated by DNA damage

A potential model to regulate YlbL and CtpA in response to DNA damage is to increase protein levels following exposure to DNA damage. Increased protease levels in response to DNA damage could promote the DNA damage checkpoint recovery when needed. To test this model, we monitored YlbL and CtpA protein levels via Western blotting over the course of the MMC recovery assay. YlbL and CtpA protein levels did not change relative to the loading control DnaN throughout the course of the experiment (Fig S4B & C). As a positive control, we performed the same experiment and monitored RecA protein levels and found that, indeed, RecA protein levels increased (Fig S4B & C), as expected because *recA* is induced as part of the SOS response (41, 42). We conclude that YlbL and CtpA protein levels are not regulated by DNA damage.

### The cell division inhibitor YneA accumulates in protease mutants

The data presented thus far led us to hypothesize that in the absence of YlbL and CtpA, a protein accumulates, resulting in inhibition of cell division (Fig 5A). To identify the accumulating protein, we performed an analysis of the entire proteome of WT and double protease mutant cell extracts. We chose to analyze the proteomes of cells after two hours of recovery, because the cell length distributions differed most between WT and the double protease mutant (Fig 4B). We found that the normalized spectral count data had similar distributions for both WT and the double mutant, which were approximately log normal (Fig S5A). We verified that the distribution of the test statistic (the difference in double mutant average and WT average) was normally distributed (Fig S5B), thus allowing a t-test to be used. We also performed a principle component analysis and found that WT replicates and double mutant replicates each clustered together (Fig S5C).

**Figure 5.**
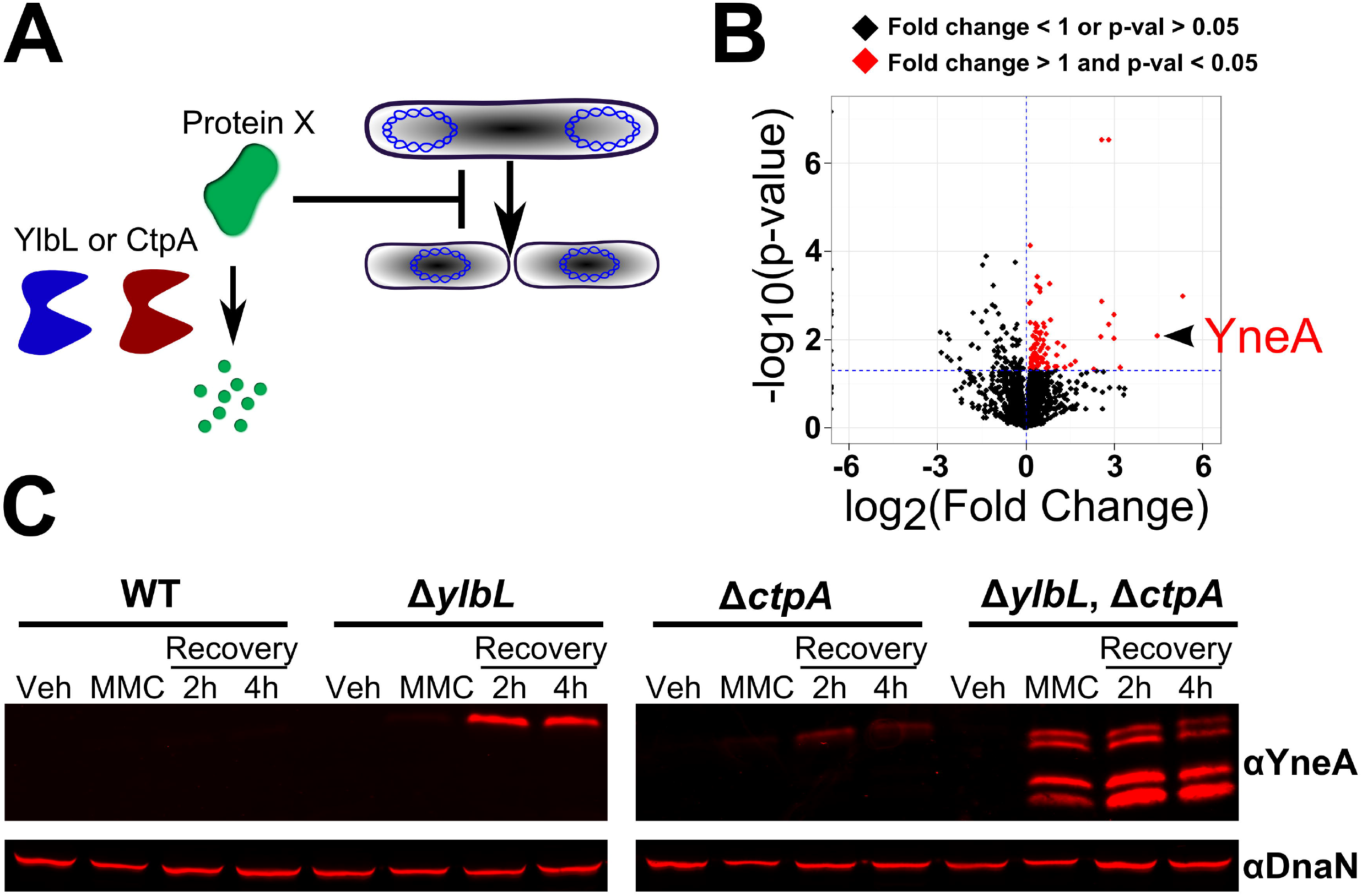
YneA accumulates in protease mutants. **(A)** A model for the function of YlbL and CtpA in regulating cell division. **(B)** Proteomics data plotted as fold change (Double Mutant/WT) vs. the p-value. Points plotted in black have a fold change less than one or a p-value greater than or equal to 0.05, and points plotted in red have a fold change greater than one and a p-value less than 0.05. **(C)** Western blot analysis of cell lysates from strains with the indicated genotypes at the indicated time points from the MMC recovery assay (see methods), using YneA or DnaN antiserum.

In total, 2329 proteins were detected, and 183 proteins were found to be differentially represented (p-value < 0.05) in the double mutant relative to WT (Table S4). Of the proteins differentially represented in the double mutant, 104 had a fold change greater than one (Fig 5B, red points). There are three major mechanisms that have been reported in *B. subtilis* to inhibit cell division: 1) Noc dependent nucleoid occlusion (43), 2) FtsL depletion (44, 45), and 3) expression of YneA (22). One possibility was that Noc protein levels were higher in the double mutant, but we observed no difference in Noc levels (Fig S5D). Another possibility was that FtsL or the protease RasP, which degrades FtsL, was affected in the protease mutant (45). We found no difference in relative protein abundance of FtsL or RasP (Fig S5D), ruling out the FtsL RasP pathway. Among the top 10 proteins that were more abundant in the double mutant was YneA, the SOS-dependent cell division inhibitor (Table S4). We asked if the enrichment of YneA was simply because it is SOS induced. We analyzed the relative abundance of several other proteins that are known to be SOS induced, including RecA, UvrA, UvrB, DinB, and YneB (41), which is in an operon with YneA (22). We found that none of these other proteins were enriched in the double mutant (Fig S5E). These results suggest that YneA accumulation is not a result of increased SOS activation, and regulation of YneA accumulation is likely to be post translational, because the protein levels of another member of the operon, YneB were unchanged. Taken together, our proteomics data suggest that YlbL and CtpA promote DNA damage checkpoint recovery through regulating YneA protein abundance.

We directly tested for YneA accumulation in protease mutants throughout the MMC recovery assay using Western blotting. YneA accumulated in all protease deletion strains after 2 hours and 4 hours of recovery, though YneA accumulation in Δ*ctpA* was slight (Fig 5C). In the double mutant, YneA accumulated in the MMC treatment condition in addition to both recovery time points (Fig 5C). In the double mutant we observed multiple YneA species, which we hypothesize to be the result of unnaturally high YneA protein levels resulting in non-specific protease activity by other proteases. With these results, we suggest that YneA is a substrate of YlbL and CtpA, both of which degrade YneA allowing for checkpoint recovery.

### *yneA* is required for DNA damage sensitivity and cell elongation phenotypes

Although accumulation of YneA fit our data well, we considered that the other proteins enriched greater than five-fold in the double mutant may have contributed to the DNA damage sensitivity phenotype. To test this, we constructed deletions of each gene in WT and the double mutant and tested for MMC sensitivity. We found that no single deletion of each of the 10 genes resulted in sensitivity to MMC (Fig S6A). In the double mutant, only deletion of *yneA* was able to rescue the sensitivity to MMC (Fig S6A). We verified that deletion of *yneA* could rescue MMC sensitivity in all protease mutant backgrounds (Fig 6A). We examined cell length in the DNA damage recovery assay. As expected, deletion of *yneA* resulted in less severe cell elongation relative to WT (compare WT in Fig 4B and Δ*yneA::loxP* in Fig 6B). In addition, deletion of *ylbL, ctpA*, or both no longer changed the cell length distribution in the absence of yneA at the two hour recovery time point (Fig 6B, 6C, and S6B). In the MMC treatment, we did observe a slight increase (p-value = 0.0004 for one-tailed Z-test) in the percentage of cells greater than 6.75 μm in the double protease deletion strain compared to WT (Fig 6C). Given that MMC sensitivity and most cell elongation in protease mutants depends on *yneA*, we hypothesized that expression of YneA alone would be sufficient to inhibit growth to a greater extent in the protease mutants. Indeed, strains lacking YlbL, CtpA or both were more sensitive to over-expression of yneA from an IPTG inducible promoter than WT (Fig 6D). Further, we show that YneA accumulated in the protease mutant strains following *yneA* ectopic expression (Fig 6E). We conclude that YneA accumulation results in severe growth inhibition in cells lacking YlbL and CtpA.

**Figure 6.**
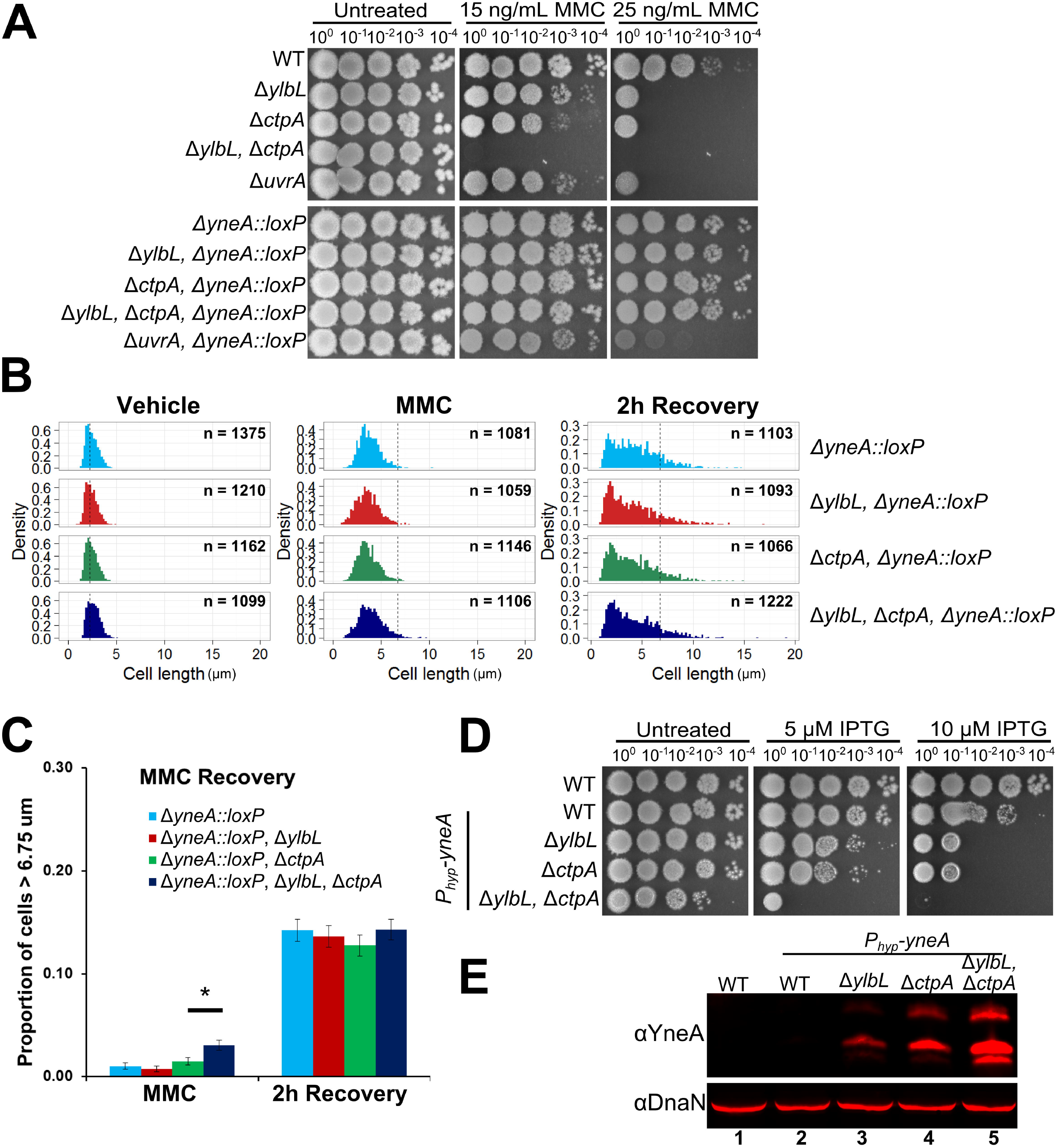
*yneA* is required for DNA damage sensitivity and cell elongation phenotypes. **(A)** Spot titer assay using the indicated genotypes and media. **(B)** Cell length distributions plotted as histograms. The number of cells scored in each distribution is indicated as “n=” and the genotype of each strain is indicated above the distributions. The dotted vertical line in the “Vehicle” distributions is at the approximate mean of 2.25 μm. The dotted vertical line in the remaining distributions is at 6.75 μm. The y-axis in all graphs is normalized by the sample size yielding the density, and the x-axis is the cell length in μm. **(C)** The proportion of cells greater than 6.75 μm from the distributions in panel B is plotted as a bar graph. The error bars represent the standard deviation, 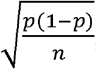, where p represents the proportion and n is the sample size). The asterisk indicates a p-value less than 0.05. **(D)** Spot-titer assay testing the effect of over production of YneA using the indicated concentration of the inducer IPTG. **(E)** Western blot analysis of cell lysates using 100 μM IPTG for YneA expression with the indicated genotypes, using the indicated antiserum.

### CtpA specifically digests YneA *in vitro*

To test the hypothesis that YneA is a direct substrate of the proteases we purified YneA, CtpA, and YlbL lacking their N-terminal transmembrane domains. We were unable to detect protease activity from YlbL using YneA, lysozyme, or casein as substrates (unpublished observations; see discussion). When purified CtpA was incubated with YneA, we observed digestion of YneA over time, but no digestion was observed using CtpA-S297A (Fig 7A). To test if CtpA activity against YneA was specific we completed the same reaction using lysozyme as a substrate and detected no activity (Fig 7B). We conclude that YneA is a direct and specific substrate of CtpA.

**Figure 7.**
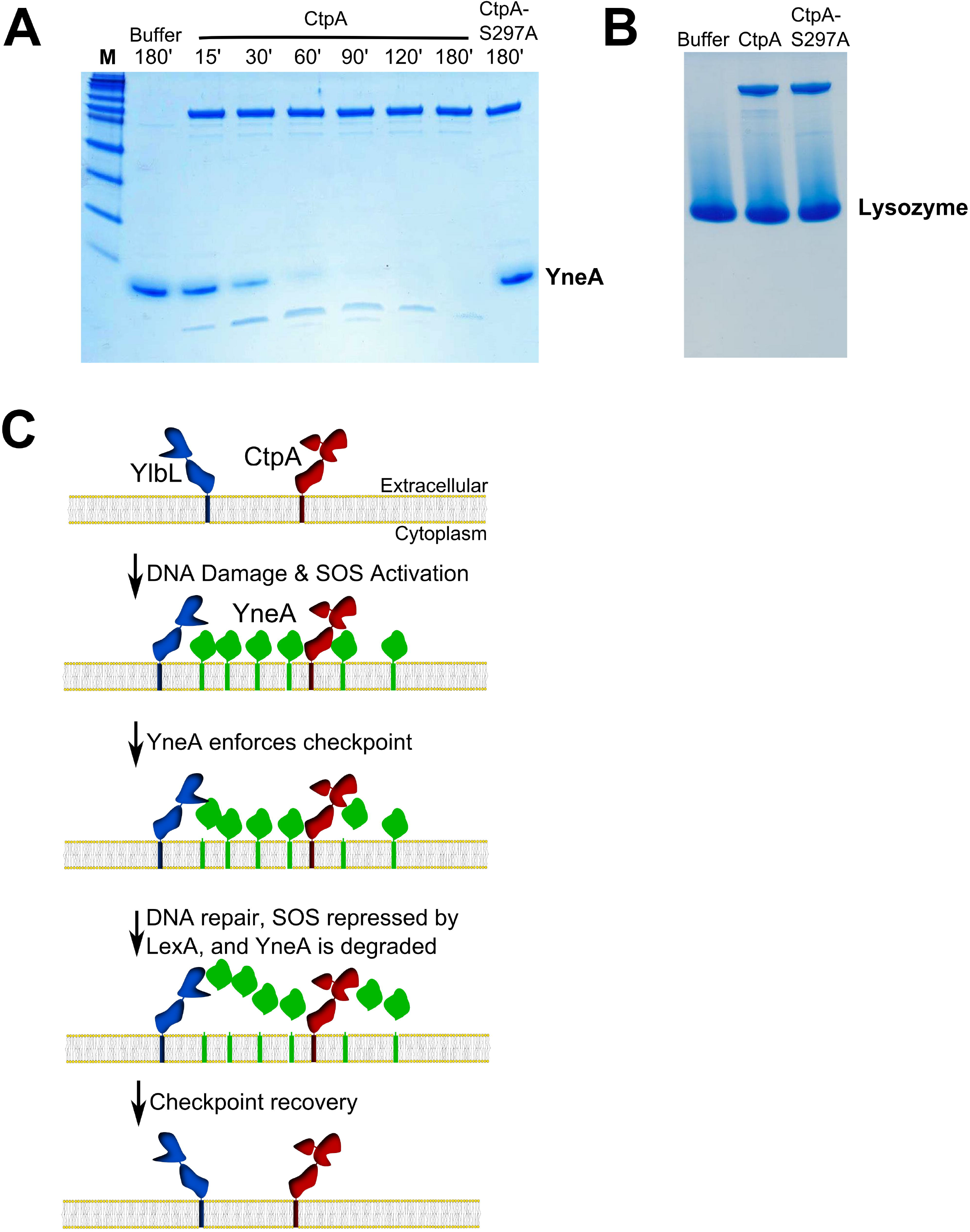
DNA damage checkpoint recovery in Bacillus subtilis. **(A)** Protease assay incubating purified CtpA or CtpA-S297A with purified YneA for the indicated time followed by SDS-PAGE and staining with coomassie blue. **(B)** Protease assay incubating purified CtpA or CtpA-S297A with commercially available lysozyme for 3 hours followed by SDS-PAGE and staining with coomassie blue. **(C)** A model for YlbL- and CtpA-dependent DNA damage checkpoint recovery. YlbL and CtpA are present as membrane proteases, and when high amounts of DNA damage are present, YneA production overwhelms both proteases resulting in delayed cell division. After DNA repair is complete and YneA expression decreases, YneA is cleared and cell division proceeds.

## Discussion

All organisms control cellular processes through regulated signaling. To regulate a cellular process a signaling pathway must have mechanisms of activation and inactivation. Many bacteria use a small membrane protein as an SOS-induced DNA damage checkpoint protein (21–24). The mechanism of checkpoint recovery, however, for organisms using membrane protein checkpoints has remained unclear. Our comprehensive study identified a dual protease mechanism of DNA damage checkpoint recovery (Fig 7C). Proteases YlbL and CtpA are constitutively present in the plasma membrane of cells even in the absence of DNA damage. After encountering DNA damage, YneA expression is induced. We hypothesize that YlbL and CtpA activities become saturated by increased YneA expression, which results in a delay of cell division. Following DNA repair, expression of YneA decreases and YlbL and CtpA clear any remaining YneA allowing cell division to resume. DNA damage checkpoints are of fundamental importance to biology, and we have discovered the pathway responsible for checkpoint inactivation and cell cycle re-entry in *B. subtilis*. With these results, we propose to rename YlbL to LmpA (<u>L</u>on <u>m</u>embrane <u>a</u>nchored <u>p</u>rotease <u>A</u>) to reflect its activity as a membrane anchored Lon protease.

Recovery from a DNA damage checkpoint is a critical process for all organisms. One theme found throughout biology is the use of multiple proteins with overlapping functions. In eukaryotes, the phosphorylation events that establish the checkpoint are removed by multiple phosphatases (46, 47). In *E. coli*, there are two cytoplasmic proteases, Lon and ClpYQ, that have been found to degrade the cell division inhibitor SulA (15, 17–19). Our study further extends the use of multiple factors in regulating checkpoint recovery to *B. subtilis*, by describing a mechanism using two extracellular proteases. In eukaryotes, multiple proteins with overlapping functions often exist due to spatial or temporal restrictions, which appears to at least partially explain the use of multiple factors in checkpoint recovery (46, 47). In *E. coli*, ClpYQ was found to be important at higher temperatures in the absence of Lon (18), again suggesting that each protease functions under specific conditions. In the case of YlbL and CtpA, however, there appears to be a shared responsibility in rich media. Deletion of each protease results in DNA damage sensitivity and the double mutant has a more severe sensitivity. In contrast, during growth in minimal media, YlbL appears to be the primary protease, as the cell elongation phenotype is more pronounced in cells lacking *ylbL*. Still it is unclear how or when each protease functions. Why isn’t one protease sufficient to degrade YneA? Do the proteases occupy distinct loci in the cell, requiring that each protease degrades a specific YneA pool? Another possibility is that protease levels are constrained by another evolutionary pressure, such as substrates unique to each protease. Thus, the cell cannot maintain the individual proteases at levels required to titrate YneA as part of the DDR, because the levels of another substrate would be too low. Another explanation is that using multiple factors is an evolutionary strategy that increases the fitness of an organism. It is clear that checkpoint recovery is crucial, because the fitness of cells lacking *ylbL* or *ctpA* is significantly decreased in the presence of DNA damage (Table S2).

Although the dual protease mechanism described here resolves an important step in the DDR, our data also reveal the complexity of the system. After we exposed cells to MMC the cells elongated. We noticed however, that not all elongation depended on *yneA* (see Fig 6B), suggesting another mechanism for cell cycle control. In *B. subtilis*, there have been reports of *yneA*-independent control of cell division following replication stress (44, 48, 49). The essential cell division component FtsL has been reported to be unstable and depletion leads to inhibition of cell division (49). Further, ftsL transcript levels were reported to decrease following replication stress independent of the SOS response (44), thus linking depletion of the unstable FtsL protein to cell division control following replication stress. A study using a replication block consisting of the Tet-repressor bound to a Tet-operator array, observed cell division inhibition independent of *yneA, noc*, and FtsL (48). Interestingly, recent studies of *Caulobacter crescentus* uncovered two cell division inhibitors that are expressed in response to DNA damage, with one SOS-dependent inhibitor and the other SOS-independent (21, 25). In *B. megaterium*, a recent study found that the transcript of *yneA* is unstable following exposure to DNA damage (50), suggesting yet another layer of regulation. No factor was identified to regulate *yneA* transcripts in the previous study, though it is possible that one of the genes of unknown function identified in our screens could regulate *yneA* mRNA. These studies highlight the complexity of regulating the DNA damage checkpoint in bacteria.

## Materials and Methods

### Bacteriological methods and chemicals

Bacterial strains, plasmids, and oligonucleotides used in this study are listed in Table S5 and the construction of strains and plasmids is detailed in the supplemental methods. All *Bacillus subtilis* strains are isogenic derivatives of PY79 (51). *Bacillus subtilis* strains were grown in LB (10 g/L NaCl, 10 g/L tryptone, and 5 g/L yeast extract) or S7_50_ minimal media with 2% glucose (1x S7_50_ salts (diluted from 10x S7_50_ salts: 104.7g/L MOPS, 13.2 g/L, ammonium sulfate, 6.8 g/L monobasic potassium phosphate, pH 7.0 adjusted with potassium hydroxide), 1x metals (diluted from 100x metals: 0.2 M MgCl_2_, 70 mM CaCl_2_, 5 mM MnCl_2_, 0.1 mM ZnCl_2_, 100 μg/mL thiamine-HCl, 2 mM HCl, 0.5 mM FeCl_3_), 0.1% potassium glutamate, 2% glucose, 40 μg/mL phenylalanine, 40 μg/mL tryptophan) at 30°C with shaking (200 rpm). Mitomycin C (MMC), methyl methane sulfonate (MMS), and phleomycin were used at the concentrations indicated in the figures. The following antibiotics were used for selection in *B. subtilis* as indicated in the method details: spectinomycin (100 μg/mL), chloramphenicol (5 μg/mL), and erythromycin (0.5 μg/mL). Selection of *Escherichia coli* (MC1061 or TOP10 cells for cloning or BL21 for protein expression) transformants was performed using the following antibiotics: spectinomycin (100 μg/mL) or kanamycin (50 μg/mL).

### Tn-seq

A transposon insertion library was constructed similar to as described (52) with modifications described in the supplemental methods. Tn-seq experiments were designed with multiple growth periods similar to a prior description (27), with a detailed description in the supplemental methods. Sequencing library construction and data analysis were performed as described previously (34, 52) with modifications described in the supplemental methods.

### Spot-titer assays

*B. subtilis* strains were struck out on LB agar and incubated at 30°C overnight. The next day, a single colony was used to inoculate a 2 mL LB culture in a 14 mL round bottom culture tube, which was incubated at 37°C on a rolling rack until OD_600_ was 0.5-1. Cultures were normalized to OD_600_ = 0.5 and serial diluted. The serial dilutions were spotted (4 μL) on the agar media indicated in the figures and the plates were incubated at 30°C overnight (16-20 hours). All spot-titer assays were performed at least twice.

### Survival assays

Survival assays using an acute treatment of mitomycin C were performed as previously described (53). Cultures were grown to an OD_600_ of about 1, and triplicate samples of 0.6 mL of an OD_600_ = 1 equivalent was taken and cells were pelleted via centrifugation: 10,000 *g* for 5 minutes at room temperature (all subsequent centrifugation steps were identical). Cells were washed with 0.6 mL 0.85% NaCl (saline) and pelleted via centrifugation. Cell pellets were resuspended in 0.6 mL saline, and 100 μL aliquots were distributed for each MMC concentration. MMC was added to each tube to yield the final concentration stated in the figure, and cells were incubated at 37°C for 30 minutes. Cells were pelleted via centrifugation to remove MMC, resuspended in saline, and a serial dilution yielding a scorable number of cells (about 30-300) was plated on LB agar to determine the surviving fraction of cells. Each experiment was performed three times in triplicate for each strain.

### Antiserum production

Purified proteins (see below for purification protocols) were submitted to Covance for antibody production using rabbits. Two rabbits were used in the 77 day protocol, and the serum with the least background was used for experiments.

### Western blotting

For YlbL, CtpA, RecA, and DnaN Western blots, a cell pellet equivalent of 1 mL OD_600_ = 1 was re-suspended in 100 μL 1x SMM buffer (0.5 M sucrose, 0.02 M maleic acid, 0.02 M MgCl_2_, adjusted to pH 6.5) containing 1 mg/mL lysozyme and 2x Roche protease inhibitors at room temperature for 1 or 2 hours. Samples were then lysed by addition of 6x SDS loading dye (0.35 M Tris, pH 6.8, 30% glycerol, 10% SDS, 0.6 M DTT, and 0.012% bromophenol blue) to 1x. Samples (12 μL) were separated via 10% SDS-PAGE, and transferred to nitrocellulose using a Trans-Blot Turbo (BioRad) according to the manufacturer’s directions. Membranes were blocked in 5% milk in TBST (25 mM Tris, pH 7.5, 150 mM NaCl, and 0.1% Tween 20) at room temperature for 1 hour or at 4°C overnight. Blocking buffer was removed, and primary antibodies were added in 2% milk in TBST (αYlbL, 1:5000 or 1:8000; αCtpA, 1:5000; αRecA, 1:4000; αDnaN, 1:4000). Primary antibody incubation was performed at room temperature for 1 hour or overnight at 4°C. Primary antibodies were removed and membranes were washed three times with TBST for 5 minutes at room temperature. Secondary antibodies (Licor; 1:15000) were added in 2% milk in TBST and incubated at room temperature for 1 hour. Membranes were washed three times as above and imaged using the Li-COR Odyssey imaging system. All Western blot experiments were performed at least twice with independent samples.

For YneA Western blots, cell pellets, 10 mL OD_600_ = 1 for MMC recovery assay and 25 mL OD_600_ = 1 for over-expression, were re-suspended in 400 or 500 μL, respectively, of sonication buffer (50 mM Tris, pH 8.0, 10 mM EDTA, 20% glycerol, 2x Roche protease inhibitors, and 5 mM PMSF), and lysed via sonication. SDS loading dye was added to 2x and samples were incubated at 100°C for 7 minutes. Samples (10 μL) were separated using 16.5% Tris-Tricine-SDS-PAGE (BioRad) and transferred to a nitrocellulose membrane using a Transblot Turbo (BioRad) according to the manufacturer’s directions. All subsequent steps were performed as above with a 1:3000 primary antibody dilution.

### Mitomycin C recovery assay

An LB agar plate grown at 30°C overnight was washed with pre-warmed S7_50_ minimal media and used to inoculate a culture of S7_50_ minimal media at an OD_600_ = 0.1. The cultures were incubated at 30°C until an OD_600_ of about 0.2 (2-2.5 hours). MMC was added to 100 ng/mL and cultures were incubated at 30°C for 2 hours. Cells were pelleted via centrifugation (4,696 *g* for 7 minutes) and the media was removed. Cell pellets were washed in an equal volume of 1x PBS, pH 7.4, and pelleted again via centrifugation as above. Cell pellets were re-suspended in an equal volume of pre-warmed S7_50_ minimal media and incubated at 30°C for four hours. Samples for microscopy and Western blot analysis were taken after the two hour MMC treatment and at two and four hours following recovery, as indicated in the figures. The vehicle control samples were treated for 2 hours with an equivalent volume of the vehicle in which MMC was suspended (25% (v/v) DMSO).

### Microscopy

A 500 μL sample from the MMC recovery assay above was taken and FM4-64 was added to 2 μg/mL and incubated at room temperature for 5 minutes. Samples were then transferred to 1% agarose pads made of 1x Spizizen’s salts. Images were captured using an Olympus BX61 microscope.

### Cell length analysis

Cells were scored for cell length using the measuring tool in ImageJ software. For each image scored, all cells that were in focus were measured. The number of cells scored for each strain/condition is stated in the figures (n=cells measured). The histograms were generated using ggplot2 in R. All scoring was done using unadjusted images. Representative images shown in the figures were modified in ImageJ by subtracting the background (rolling ball radius method) and adjusting the brightness and contrast. Any adjustments made were applied to the entire image.

### Proteomics experimental details

Samples (5 mL OD_600_ = 1) were harvested from cultures grown as described in the MMC recovery assay section at 2 hours recovery via centrifugation: 4,696 *g* at room temperature for 10 minutes. Samples were washed twice with 500 μL 1x PBS, pH 7.4 and pelleted via centrifugation: 10,000 *g* at room temperature for 5 minutes. Samples were frozen in liquid nitrogen and stored at −80°C. Samples were submitted for mass-spectrometry analysis to MS Bioworks. Further sample processing and data analysis was performed by MS Bioworks as described in the supplemental methods.

### YneA and lysozyme digestion assays

YneA digestion reactions were prepared as a 20 μL reaction in 20 mM Tris pH 7.5, 20 mM NaCl, and 20% glycerol containing 150 μM YneA, and 2 μM CtpA. Reactions were incubated at 30°C for the time indicated in the figure. Reactions were stopped by addition of 6x SDS-dye to 1x and incubating at 100°C for 5 minutes. Reaction products were separated via 16.5% Tris-Tricine SDS-PAGE. Proteins were detected by staining with coomassie blue. Lysozyme digestion assays were performed as for YneA using 2 mg/mL lysozyme and reactions were incubated at 30°C for 3 hours.

## Acknowledgements

We would like to thank members of the Simmons lab for thoughtful discussions throughout this work. We would like to thank Dr. Jayakrishnan Nandakumar for the generous gift of plasmids for protein overexpression, and Dr. David Rudner for plasmid pDR110. This work was supported by NIH grant R01 GM107312 to L.A.S. J.W.S. was supported in part by NIH T32 GM007544 and Associate Professor Funds from the College of Literature, Science and the Arts at the University of Michigan. P.E.B. was supported by a pre-doctoral fellowship from the National Science Foundation (DGE 1256260).

## Author contributions

Experiments were designed by P.E.B. and L.A.S. Experiments were performed by P.E.B. and Z.W.S. Data were analyzed by P.E.B., Z.W.S., J.W.S., and L.A.S. The paper was written by P.E.B. and L.A.S.

---

**Figure S1 Tn-seq data analysis. (A)** Sequencing read distributions for transposon (Tn) insertion locations containing greater than 0 reads for each replicate of the library samples (left) or the starter culture samples (right) from the MMC experiment. The y-axis is the frequency of Tn sites, and the x-axis is the log10 of sequencing reads. The dotted vertical line is drawn at log_10_(100). **(B)** Sequencing read distributions are plotted for the indicated samples as in panel A, except the replicates were summed prior to plotting. The library and starter cultures are the same in the Ctrl and MMC plots; the library and starter cultures are the same in the MMS and Phleo plots, which are shown twice in each case to allow direct comparison. The dotted vertical line is drawn at log_10_(350). **(C)** Relative fitness distributions are plotted for Tn insertions with more than 10 sequencing reads in the control samples. The y-axis is the frequency of Tn sites and the x-axis is the relative fitness. The dotted vertical line is drawn at 1.0. All three growth periods are plotted for the indicated experiments.

**Figure S2 YlbL and CtpA catalytic residue identification. (A)** The Lon protease domain of YlbL was aligned to LonA and LonB from *B. subtilis* and Lon from *E. coli*. The alignments show that YlbL contains the conserved catalytic dyad of Lon proteases consisting of serine 234 and lysine 279. **(B)** The S41 protease domain of CtpA was aligned to CtpB from *B. subtilis* and Prc from *E. coli*. The alignments showed that CtpA contains a conserved catalytic triad consisting of serine 297, lysine 322, and glutamine 326.

**Figure S3 Disruption of *ylbK* results in a polar effect on *ylbL***. **(A)** Spot titer assay using the indicated genotypes and media. **(B)** Schematic of *ylbK* and *ylbL* loci, with the putative ribosome binding site, proposed to control *ylbL* translation, labeled in yellow. **(C)** Western blot analysis of cell lysates of the indicated genotypes using YlbL or DnaN antiserum.

**Figure S4 YlbL and CtpA levels are not regulated by DNA damage. (A)** MMC survival assay using strains with the indicated genotypes to test if MMC sensitivity is caused by cell death. The concentration of MMC used during a 30 minute incubation is listed on the x-axis, and the y-axis is the percent of cells surviving the treatment relative to the no treatment (0 ng/mL) condition. Each point is the average of three technical replicates from three individual experiments (n=9), and the error bars represent the standard error of the mean. **(B)** Representative Western blot analysis of cell lysates throughout the MMC recovery assay using YlbL, CtpA, DnaN, or RecA antiserum. **(C)** Quantification of Western blot data plotted as a bar graph. The bars represent the average from three experiments (YlbL, CtpA, and DnaN) or two experiments (RecA), and the error bars are the standard deviation (YlbL and CtpA) or the range (RecA) of the measurements. The y-axis is the relative protein levels, which is the indicated protein level normalized to the loading control, DnaN, and the no treatment measurement.

**Figure S5 YneA accumulates in protease mutants. (A)** The averages of the normalized spectral counts are plotted as histograms for WT (blue) and Δ*ylbL*, Δ*ctpA* double mutant (DM; red). The y-axis is the count and the x-axis is the log_10_(normalized spectral counts for the average of three replicates). **(B)** The distribution of the test statistic (WT average - DM average) is plotted as a histogram. **(C)** A principle component analysis was performed using the normalized spectral counts from WT (blue) and DM (red) samples using the “prcomp” function in R. The first two coordinates are plotted as the x- and y-axes, respectively. **(D & E)** The average relative protein levels (WT/DM) from the proteomics dataset are plotted for the indicated proteins, and the error bars represent the standard deviation. The inset in panel E shows a closer look around one for clarity.

**Figure S6 *yneA* is required for DNA damage sensitivity and cell elongation phenotypes. (A)** Strains with the indicated genotypes (plates at left) were struck onto the indicated media (column labels) and incubated at 30°C overnight. Deletion of *yneA* suppresses the Δ*ylbL*, Δ*ctpA* double mutant (DM) MMC sensitivity phenotype. **(B)** Representative micrographs of cells stained with FM4-64 from the indicated genotypes at the indicated time points in the MMC recovery assay. The scale bar is 5 μm.

**Table S1 Tn-seq data collected**. OD_600_ measurements, incubation times, viable cell counts, growth rates estimated based on viable cell counts, number of generations estimated based on viable cell counts and incubation times, sequencing sample IDs, sequencing reads, reads mapped, and the number of Tn insertions with more than 10 reads for each sample are presented.

**Table S2 Tn-seq relative fitness lists**. The relative fitness values for each gene with sufficient data in all three growth periods for all three Tn-seq experiments are presented along with the adjusted p-value (BH method; see STAR methods). The gene names or locus tags are listed, and intergenic regions are annotated as “ig” with a number.

**Table S3 Tn-seq list overlaps. (First tab)** The genes with the lowest relative fitness with an adjusted p-value less than 0.01 are listed for each experiment. **(Second tab)** The genes overlapping in all three Tn-seq experiments as well as genes overlapping in pairwise comparisons of the three Tn-seq experiments are presented.

**Table S4 Proteomics data set**. The proteomics data for the 183 proteins identified as having significantly different levels (p-value < 0.05) in the protease double mutant relative to the control are presented.

**Table S5 Oligonucleotides, plasmids, and strains used in this study**.

## References

1. Jackson SP, Bartek J. The DNA-damage response in human biology and disease. Nature. 2009;461(7267):1071–8.

2. Blanpain C, Mohrin M, Sotiropoulou PA, Passegue E. DNA-damage response in tissue-specific and cancer stem cells. Cell stem cell. 2011;8(1):16–29.

3. Kreuzer KN. DNA damage responses in prokaryotes: regulating gene expression, modulating growth patterns, and manipulating replication forks. Cold Spring Harb Perspect Biol. 2013;5(11):a012674.

4. Michel B. After 30 years of study, the bacterial SOS response still surprises us. PLoS biology. 2005;3(7):e255.

5. Baharoglu Z, Mazel D. SOS, the formidable strategy of bacteria against aggressions. FEMS microbiology reviews. 2014;38(6):1126–45.

6. Friedberg EC, Walker GC, Siede W, Wood RD, Schultz RA, Ellenberger T. DNA Repair and Mutagenesis. 2nd ed. Washington, D.C.: ASM Press; 2006.

7. Sancar A, Lindsey-Boltz LA, Unsal-Kacmaz K, Linn S. Molecular mechanisms of mammalian DNA repair and the DNA damage checkpoints. Annual review of biochemistry. 2004;73:39–85.

8. Ciccia A, Elledge SJ. The DNA damage response: making it safe to play with knives. Mol Cell. 2010;40(2):179–204.

9. Huisman O, D’Ari R. An inducible DNA replication-cell division coupling mechanism in E. coli. Nature. 1981;290(5809):797–9.

10. Huisman O, D’Ari R, Gottesman S. Cell-division control in Escherichia coli: specific induction of the SOS function SfiA protein is sufficient to block septation. Proceedings of the National Academy of Sciences of the United States of America. 1984;81(14):4490–4.

11. Bi E, Lutkenhaus J. Cell division inhibitors SulA and MinCD prevent formation of the FtsZ ring. Journal of bacteriology. 1993;175(4):1118–25.

12. Huang J, Cao C, Lutkenhaus J. Interaction between FtsZ and inhibitors of cell division. Journal of bacteriology. 1996;178(17):5080–5.

13. Mukherjee A, Cao C, Lutkenhaus J. Inhibition of FtsZ polymerization by SulA, an inhibitor of septation in Escherichia coli. Proceedings of the National Academy of Sciences of the United States of America. 1998;95(6):2885–90.

14. Trusca D, Scott S, Thompson C, Bramhill D. Bacterial SOS checkpoint protein SulA inhibits polymerization of purified FtsZ cell division protein. Journal of bacteriology. 1998;180(15):3946–53.

15. Mizusawa S, Gottesman S. Protein degradation in Escherichia coli: the lon gene controls the stability of sulA protein. Proceedings of the National Academy of Sciences of the United States of America. 1983;80(2):358–62.

16. Sonezaki S, Ishii Y, Okita K, Sugino T, Kondo A, Kato Y. Overproduction and purification of SulA fusion protein in Escherichia coli and its degradation by Lon protease in vitro. Applied microbiology and biotechnology. 1995;43(2):304–9.

17. Canceill D, Dervyn E, Huisman O. Proteolysis and modulation of the activity of the cell division inhibitor SulA in Escherichia coli lon mutants. Journal of bacteriology. 1990;172(12):7297–300.

18. Wu WF, Zhou Y, Gottesman S. Redundant in vivo proteolytic activities of Escherichia coli Lon and the ClpYQ (HslUV) protease. Journal of bacteriology. 1999;181(12):3681–7.

19. Seong IS, Oh JY, Yoo SJ, Seol JH, Chung CH. ATP-dependent degradation of SulA, a cell division inhibitor, by the HslVU protease in Escherichia coli. FEBS letters. 1999;456(1):211–4.

20. Kanemori M, Yanagi H, Yura T. The ATP-dependent HslVU/ClpQY protease participates in turnover of cell division inhibitor SulA in Escherichia coli. Journal of bacteriology. 1999;181(12):3674–80.

21. Modell JW, Hopkins AC, Laub MT. A DNA damage checkpoint in Caulobacter crescentus inhibits cell division through a direct interaction with FtsW. Genes & development. 2011;25(12):1328–43.

22. Kawai Y, Moriya S, Ogasawara N. Identification of a protein, YneA, responsible for cell division suppression during the SOS response in Bacillus subtilis. Molecular microbiology. 2003;47(4):1113–22.

23. Ogino H, Teramoto H, Inui M, Yukawa H. DivS, a novel SOS-inducible cell-division suppressor in Corynebacterium glutamicum. Molecular microbiology. 2008;67(3):597–608.

24. Chauhan A, Lofton H, Maloney E, Moore J, Fol M, Madiraju MV, et al. Interference of Mycobacterium tuberculosis cell division by Rv2719c, a cell wall hydrolase. Molecular microbiology. 2006;62(1):132–47.

25. Modell JW, Kambara TK, Perchuk BS, Laub MT. A DNA damage-induced, SOS-independent checkpoint regulates cell division in Caulobacter crescentus. PLoS biology. 2014;12(10):e1001977.

26. Mo AH, Burkholder WF. YneA, an SOS-induced inhibitor of cell division in Bacillus subtilis, is regulated posttranslationally and requires the transmembrane region for activity. Journal of bacteriology. 2010;192(12):3159–73.

27. Byrne RT, Chen SH, Wood EA, Cabot EL, Cox MM. Escherichia coli genes and pathways involved in surviving extreme exposure to ionizing radiation. Journal of bacteriology. 2014;196(20):3534–45.

28. Iyer VN, Szybalski W. A molecular mechanism of mitomycin action: Linking of complementary DNA strands. Proceedings of the National Academy of Sciences of the United States of America. 1963;50:355–62.

29. Noll DM, Mason TM, Miller PS. Formation and repair of interstrand cross-links in DNA. Chemical reviews. 2006;106(2):277–301.

30. Sedgwick B. Repairing DNA-methylation damage. Nature reviews Molecular cell biology. 2004;5(2):148–57.

31. Reiter H, Milewskiy M, Kelley P. Mode of action of phleomycin on Bacillus subtilis. Journal of bacteriology. 1972;111(2):586–92.

32. Kross J, Henner WD, Hecht SM, Haseltine WA. Specificity of deoxyribonucleic acid cleavage by bleomycin, phleomycin, and tallysomycin. Biochemistry. 1982;21(18):4310–8.

33. Robinson DG, Chen W, Storey JD, Gresham D. Design and analysis of Bar-seq experiments. G3 (Bethesda, Md). 2014;4(1):11–8.

34. van Opijnen T, Bodi KL, Camilli A. Tn-seq: high-throughput parallel sequencing for fitness and genetic interaction studies in microorganisms. Nature methods. 2009;6(10):767–72.

35. van Opijnen T, Camilli A. Transposon insertion sequencing: a new tool for systems-level analysis of microorganisms. Nature reviews Microbiology. 2013;11(7):435–42.

36. Benjamini Y, Hochberg Y. Controlling the false discovery rate: a practical and powerful approach to multiple testing. J R Stat Soc Ser B-Methodol. 1995;57(1):289–300.

37. Lenhart JS, Schroeder JW, Walsh BW, Simmons LA. DNA repair and genome maintenance in Bacillus subtilis. Microbiology and molecular biology reviews : MMBR. 2012;76(3):530–64.

38. Sancar A. DNA excision repair. Annual review of biochemistry. 1996;65:43–81.

39. Friedman BM, Yasbin RE. The genetics and specificity of the constitutive excision repair system of Bacillus subtilis. Molecular & general genetics : MGG. 1983;190(3):481–6.

40. Walsh BW, Bolz SA, Wessel SR, Schroeder JW, Keck JL, Simmons LA. RecD2 helicase limits replication fork stress in Bacillus subtilis. Journal of bacteriology. 2014;196(7):1359–68.

41. Au N, Kuester-Schoeck E, Mandava V, Bothwell LE, Canny SP, Chachu K, et al. Genetic composition of the Bacillus subtilis SOS system. Journal of bacteriology. 2005;187(22):7655–66.

42. Goranov AI, Kuester-Schoeck E, Wang JD, Grossman AD. Characterization of the global transcriptional responses to different types of DNA damage and disruption of replication in Bacillus subtilis. Journal of bacteriology. 2006;188(15): 5595–605.

43. Wu LJ, Errington J. Coordination of cell division and chromosome segregation by a nucleoid occlusion protein in Bacillus subtilis. Cell. 2004;117(7):915–25.

44. Goranov AI, Katz L, Breier AM, Burge CB, Grossman AD. A transcriptional response to replication status mediated by the conserved bacterial replication protein DnaA. Proceedings of the National Academy of Sciences of the United States of America. 2005;102(36):12932–7.

45. Bramkamp M, Weston L, Daniel RA, Errington J. Regulated intramembrane proteolysis of FtsL protein and the control of cell division in Bacillus subtilis. Molecular microbiology. 2006;62(2):580–91.

46. Shaltiel IA, Krenning L, Bruinsma W, Medema RH. The same, only different - DNA damage checkpoints and their reversal throughout the cell cycle. Journal of cell science. 2015;128(4):607–20.

47. Wang H, Zhang X, Teng L, Legerski RJ. DNA damage checkpoint recovery and cancer development. Experimental cell research. 2015;334(2):350–8.

48. Bernard R, Marquis KA, Rudner DZ. Nucleoid occlusion prevents cell division during replication fork arrest in Bacillus subtilis. Molecular microbiology. 2010;78(4):866–82.

49. Daniel RA, Harry EJ, Katis VL, Wake RG, Errington J. Characterization of the essential cell division gene ftsL(yIID) of Bacillus subtilis and its role in the assembly of the division apparatus. Molecular microbiology. 1998;29(2):593–604.

50. Buchholz M, Nahrstedt H, Pillukat MH, Deppe V, Meinhardt F. yneA mRNA instability is involved in temporary inhibition of cell division during the SOS response of Bacillus megaterium. Microbiology (Reading, England). 2013;159(Pt 8):1564–74.

51. Youngman P, Perkins JB, Losick R. Construction of a cloning site near one end of TN917 into which foreign DNA may be inserted without affecting transposition in Bacillus subtilis or expression of the transposon-bourne ERM gene. Plasmid. 1984;12(1):1–9.

52. Johnson CM, Grossman AD. Identification of host genes that affect acquisition of an integrative and conjugative element in Bacillus subtilis. Molecular microbiology. 2014;93(6):1284–301.

53. Burby PE, Simmons LA. MutS2 Promotes Homologous Recombination in Bacillus subtilis. Journal of bacteriology. 2017;199(2).

54. Sanchez H, Kidane D, Castillo Cozar M, Graumann PL, Alonso JC. Recruitment of Bacillus subtilis RecN to DNA double-strand breaks in the absence of DNA end processing. Journal of bacteriology. 2006;188(2):353–60.

55. Mascarenhas J, Sanchez H, Tadesse S, Kidane D, Krisnamurthy M, Alonso JC, et al. Bacillus subtilis SbcC protein plays an important role in DNA inter-strand cross-link repair. BMC Mol Biol. 2006;7:15.

56. Cardenas PP, Carrasco B, Defeu Soufo C, Cesar CE, Herr K, Kaufenstein M, et al. RecX facilitates homologous recombination by modulating RecA activities. PLoS Genet. 2012;8(12):e1003126.

